# CAZyme3D: a database of 3D structures for carbohydrate-active enzymes

**DOI:** 10.1101/2024.12.27.630555

**Authors:** N.R. Siva Shanmugam, Yanbin Yin

## Abstract

CAZymes (<u>C</u>arbohydrate <u>A</u>ctive En<u>Zymes</u>) degrade, synthesize, and modify all complex carbohydrates on Earth. CAZymes are extremely important to research in human health, nutrition, gut microbiome, bioenergy, plant disease, and global carbon recycling. Current CAZyme annotation tools are all based on sequence similarity. A more powerful approach is to detect protein structural similarity between query proteins and known CAZymes indicative of distant homology. Here, we developed CAZyme3D (https://pro.unl.edu/CAZyme3D/) to fill the research gap that no dedicated 3D structure databases are currently available for CAZymes. CAZyme3D contains a total of 870,740 AlphaFold predicted 3D structures (named Whole dataset). A subset of CAZymes 3D structures from 188,574 nonredundant sequences (named ID50 dataset) were subject to structural similarity-based clustering analyses. Such clustering allowed us to organize all CAZyme structures using a hierarchical classification, which includes existing levels defined by the CAZy database (class, clan, family, subfamily) and newly defined levels (subclasses, structural cluster [SC] groups, and SCs). The inter-family structural clustering successfully grouped CAZy families and clans with the same structural folds in the same subclasses. The intra-family structural clustering classified structurally similar CAZymes into SCs, which were further classified into SC groups. SCs and SC groups differed from sequence similarity-based CAZy subfamilies. With CAZyme structures as the search database, we created job submission pages, where users can submit query protein sequences or PDB structures for a structural similarity search. CAZyme3D will be a useful new tool to assist the discovery of novel CAZymes by providing a comprehensive database of CAZyme 3D structures.

## Introduction

CAZymes (<u>C</u>arbohydrate <u>A</u>ctive En<u>Zymes</u>) are enzymes that primarily target glycosidic linkages to degrade, synthesize, or modify all carbohydrates on Earth [1]. CAZymes are very abundant in plants and plant-associated microbes [2]. For example, the digestive system of humans and other animals is a carbohydrate rich environment with a very high diversity of carbohydrate-degrading bacteria. The combined CAZyme repertoire of the human gut microbiome numbers into tens of thousands of added genes [3]. CAZymes are extremely important for human health, nutrition, gut microbiome, bioenergy, plant disease, and global carbon recycling.

To enable automated CAZyme annotation, we developed multiple databases (dbCAN-seq [2, 4]), dbCAN-PUL [5], dbCAN-sub [6, 7]), web servers (dbCAN [7-9], pHMM-tree [10]), and standalone software packages (run_dbcan [9], CGC-Finder [7], eCAMI [6], dbCAN-profiler [11]). All current CAZyme annotation tools (e.g., run_dbcan, eCAMI, and CUPP [12]) assign query proteins to established CAZyme families based on sequence similarity (conserved functional domains or discrete peptide motifs). Therefore, they are unable to predict entirely new CAZyme families. Protein structures are more conserved than sequences in evolution, and protein structural similarity search is a more powerful approach to detect distant homology between functionally uncharacterized proteins and characterized CAZymes. However, less than 10,000 CAZymes have experimentally solved PDB structures, in contrast to 2.8 million sequences in the expert curated CAZy database (only 0.3% with 3D structures) [13].

Since 2021, major breakthroughs have happened in artificial intelligence (AI), enabling accurate protein structure predictions, rapid structure alignments, and scalable structure clustering for large protein sequence databases like UniProt [14] and MGnify [15]. For example, AlphaFold2 [16], ESMFold [17], TM-Vec [18], Foldseek [19], and other highly innovative software tools [20, 21] are released, breaking the technical barriers that prevented large-scale protein 3D structure predictions and comparisons that were impossible five years ago. However, in the CAZyme research field, no web resources are available to collect and analyze AlphaFold2 predicted CAZyme 3D structures, and no tools allow CAZyme annotation using the more sensitive structural similarity search methods (e.g., TM-Vec and Foldseek). To fill the research gap, we developed CAZyme3D (https://pro.unl.edu/CAZyme3D/) as the first online database dedicated for high quality CAZyme 3D structures predicted by AlphaFold2.

The CAZy database (www.cazy.org) provides PDB links to less than 10,000 structures. The CAZyme3D presented here contains over 870,000 AlphaFold2 predicted 3D structures. We provide a structural similarity search function on our website to allow users compare their own protein sequences or structures against the structures in CAZyme3D. This enables a more sensitive discovery of entirely novel CAZyme families that are unable to detect by the sequence similarity search, the limitation of all current CAZyme annotation tools. Furthermore, various visualization and analysis functions are developed to facilitate the comparisons of intra-family and inter-family structural similarities of CAZymes. CAZyme3D will be updated yearly to include AlphaFold2 structures of new CAZymes.

## Materials and Methods

### Data collection

All proteins in CAZyme3D have been annotated as CAZymes by the expert curated CAZy database [13]. All proteins have GenBank accession numbers. Their protein sequences (2.8 million) are available at https://bcb.unl.edu/dbCAN2/download/CAZyDB.07262023.fa. To remove sequence redundancy, we processed the 2.8 million CAZymes using CD-HIT [22] (-c 0.5) and obtained 188,574 sequences with <50% sequence identity among each other. This non-redundant dataset is named CAZyID50 (**Figure 1a**), with each sequence representing a cluster of sequences that share ≥ 50% sequence identity with the representative sequence.

**Figure 1:**
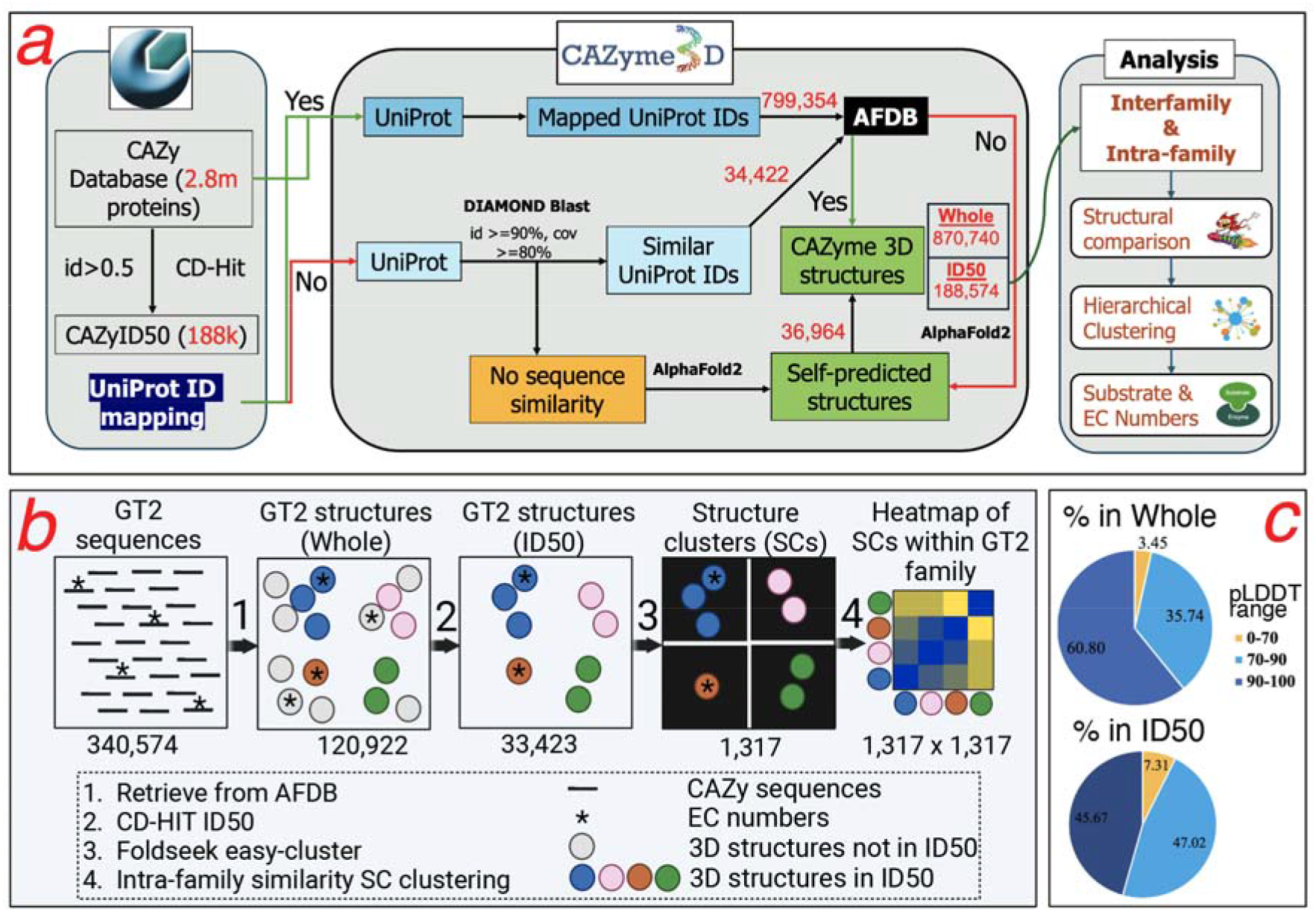
Overview of the construction of CAZyme3D database and data analysis. (**a**) The pipeline to generate CAZyme 3D structures, functional information, and comparisons for intra-family and inter-family analysis. (**b**) The illustration of intra-family analysis of CAZyme structures using GT2 family as an example. Numbers below the squares are the counts of sequences, structures, SCs, and heatmap rows x columns, respectively. (**c**) pLDDT (predicted local-distance difference test) score distribution of structures in the Whole and ID50 datasets of CAZyme3D.

For the 2.8 million CAZy proteins, we searched their GenBank IDs in the UniProt ID mapping file (https://ftp.uniprot.org/pub/databases/uniprot/current_release/knowledgebase/idmapping/) to obtain their UniProt IDs. With the UniProt IDs, we were able to retrieve 3D structures in the PDB format for 799,354 CAZymes from the AlphaFold Structure Database (AFDB) [23] (**Figure 1a**).

Among these 799,354 CAZymes, 117,188 are representative sequences in the non-redundant CAZyID50 dataset (**Figure 1a**). For the rest of the CAZymes (71,386) in CAZyID50, we searched them against the UniProt database using DIAMOND [24]. For 34,422 of them, we found their best match protein in AFDB that shares sequence identity ≥ 90% and coverage ≥ 80% and obtained their AlphaFold 3D structures. For the remaining 36,964 CAZymes, we predicted their 3D structures using AlphaFold2 via ColabFold v1.5.3 [25].

Together, we obtained a total of 870,740 CAZyme 3D structures, forming the “Whole” CAZyme3D dataset (**Figure 1a**). The “ID50” dataset is a subset of the “Whole” dataset, consisting of 188,574 CAZyme 3D structures (117,188 from UniProt ID mapping, 34,422 from DIAMOND search, and 36,964 self-predicted) in CAZyID50.

### Structural clustering

According to the CAZy annotation, CAZymes are classified into CAZy families. For each CAZy family, we took all 3D structures of the ID50 dataset and clustered them using the Foldseek easy-cluster command (alignment coverage -c 0.9). The idea was to generate structural clusters (SCs, equivalent to subfamilies) within each family (see **Figure 1b** for GT2 family as an example). As such, we organized all structures of each family in a hierarchical classification: each family (e.g., GT2) has multiple SCs, each SC contains multiple ID50 structures sharing structural similarity, and each ID50 structure represents multiple structures sharing > 50% sequence similarity in the Whole dataset. For CBM structures, we extracted CBM domains (predicted by dbCAN [9]) and only used the CBM domain structures to generate CBM SCs. For other classes, we used the full-length protein structures for clustering.

### Intra-family and inter-family structural similarity heatmaps

Furthermore, we calculated the average TM-score (template modelling score, a measure of similarity between two structures) for each pair of SCs within each CAZy family (GT2 as an example in **Figure 1b**). For example, SC1 has *m* structures, and SC2 has *n* structures. The TM-score for SC1-SC2 is averaged from *m* x *n* TM-scores calculated by Foldseek. Then, hierarchical clustering was applied to all SCs based on their pair-wise TM-scores and plotted as a heatmap. For each CAZy family, we provided on our website an interactive heatmap as the intra-family structural similarity comparison result for all SCs.

CAZy classifies ∼500 CAZyme families into six classes: glycosyltransferases (GTs), glycoside hydrolases (GHs), polysaccharide lyases (PLs), carbohydrate esterases (CEs), enzymes of auxiliary activities (AAs), and carbohydrate binding modules (CBMs). Following the procedure for the intra-family structural similarity comparison among SCs, we also performed the inter-family structural similarity comparisons and clustering. Specifically, the structural similarities between each pair of CAZyme families within the same class were calculated. For example, the GH class has 184 CAZyme families. We calculated the average TM-score for each pair of GH families (e.g., GH5 vs. GH13) and plotted the hierarchical clustering of all 184 GH families as a heatmap based on their pair-wise TM-scores. For each CAZy class, we provided on our website an interactive heatmap as the inter-family structural similarity comparison result for all families of the class.

### Substrate and EC mapping

A small fraction (∼10,000 out of 2.8 million) of CAZymes are experimentally characterized with EC (enzyme commission) numbers so that their glycan substrates are known. Using the CAZyme EC and substrate mapping table (https://bcb.unl.edu/dbCAN2/download/dbsub_data/fam-substrate-mapping-08252022.tsv) from dbCAN-sub [7], we found 8,163 CAZymes have EC numbers and are mapped to glycan substrates in CAZyme3D. CAZymes and their EC and substrate information are provided in the SC (subfamily) pages of CAZyme3D, which will help infer more specific functions for each SC (subfamily).

### Web development

The CAZyme3D website was developed using PHP and Python for request handling and server-side routing. For the user interface layout, we employed AdminLTE-2.3.11 (https://github.com/heliofidalgo/AdminLTE-2.3.11) based on the Bootstrap 3 (https://getbootstrap.com/docs/3.3/) framework and custom stylesheets to complement Bootstrap’s base styling. We used a MySQL server version (8.0.35) for data storage and PHP codes to interact with our database. Interactive heatmaps and clustered maps were visualized using the Plotly.js (v2.35.0) (https://plotly.com/).

### Structural similarity search web service

To allow users to search their query proteins against CAZyme3D, we constructed local searchable databases to enable a structural similarity search using sequence and structure inputs. Two structural similarity search tools were used.

The first tool is TM-Vec [18]. Users can submit protein sequences in FASTA format as query for a structure-aware search against our pre-computed TM-Vec databases. We constructed five TM-Vec databases with the *tmvec-build-database* command: (1) CAZy Characterized (10,977 CAZy proteins with EC numbers), (2) CAZy ID50 (188,574 CAZy proteins with structures), (3) CAZy Whole (2.8 million CAZy proteins), (4) CATH [26], and (5) PDB [27].

The second tool is Foldseek [19]. Users can submit a sequence in FASTA format or a 3D structure in PDB format for a structural similarity search against our pre-computed Foldseek databases. For a PDB query search, we constructed two Foldseek databases with the *createdb* command: (1) CAZyme3D Whole (870,740 structures) and (2) CAZyme3D-ID50 (188,574 structures).

### Keyword search and data browse options

A keyword search function is provided for users to search for CAZyme structures by GenBank and UniProt IDs, CAZy families and subfamilies, or filter by the predicted local-distance difference test (pLDDT) score (a measure of per-residue accuracy of the predicted structure).

At the homepage of our website, users can browse by taxonomy and by CAZy family. The result page is shown as a table with columns including CAZy family and subfamily, taxonomy, CAZyID50 ID and representative GenBank ID, SC ID, UniProt ID, and pLDDT score. Users can also browse by inter-family and intra-family comparisons, and the result page shows the interactive heatmap, which can be downloaded with different formats (PDF, SVG, PNG, HTML).

From the browse or keyword search result tables, users can be directed to the individual protein structure page. For each CAZyme of the Whole CAZyme3D dataset (870,740 structures), the protein page includes the following information: (i) Basic Information such as CAZy family, sequence length, UniProt ID, pLDDT score, CAZyID50 ID and representative sequence ID, (ii) Taxonomy with NCBI Taxonomy ID and lineage, (iii) CAZyme sequence and 3D structure visualization, (iv) Predicted carbohydrate binding sites by CAPSIF [28], (v) Predicted dbCAN domains, and (vi) Sequence similarity to the ID50 representative (DIAMOND result). The NGL Viewer web application (https://github.com/nglviewer/ngl) [29] is used to visualize the structures, domains, and binding sites.

## Results

### Database content: most CAZymes in CAZyme3D have highly confident AFDB structures

CAZyme3D contains a total of 870,740 CAZyme structures (the Whole dataset in **Figure 1a**). A majority of them (799,354 or 92%) are directly retrieved from AFDB by mapping CAZy GenBank IDs to UniProt IDs and then to AFDB (see Methods). A subset of the Whole dataset, designated as the ID50 dataset, contains 188,574 structures. Protein sequences of the ID50 structures are representative sequences of the Whole dataset after removing sequence redundancy with <50% sequence identity. Regarding the quality of CAZyme 3D structures, the pLDDT distribution plot shows that 92.69% and 96.54% of CAZymes have highly confident (>70 pLDDT score) AlphaFold predicted structures in the ID50 dataset and the Whole dataset, respectively (**Figure 1c**). Regarding the taxonomical distribution, 77.8% of the 870,740 CAZyme structures (Whole dataset) are from Bacteria, followed by Eukaryota (19.8%), Archaea (1.9%), Viruses (0.3%), and Unassigned (0.2%).

### Inter-family comparison: structural similarity guided clustering of CAZyme families

The 870,740 CAZymes belong to 461 protein families of 6 classes (AAs, CEs, PLs, GHs, GTs, and CBMs) according to the CAZy classification. CAZy defines each family based on sequence similarity. Different families of the same class share no or very low sequence similarity but may have structural similarity indicating a similar evolutionary origin. CAZy groups 76 out of 184 GH families into 20 clans, where a clan is a level of classification between family and class. Families of a same clan are known to share the similar structural fold, e.g., the largest clan GH-A contains 28 GH families having the (β/α)_8_ barrel fold.

To study how all families of each CAZyme class are structurally related, we have performed a structural similarity guided hierarchical clustering of families within each class and plotted the clustering as a heatmap (**Figure 2a** for GH class, see Methods). The average TM-scores (*alntmscore*) of all family pairs within each of the six classes are: AA: 0.27, CE: 0.28, PL: 0.25, GH: 0.23, GT: 0.28, and CBM: 0.23. Even though these average inter-family TM-scores are low, the hierarchical clustering reveals high structural similarities between different families (**Figure 2a**). For example, the GH class heatmap shows that 36 families with the (α/α)_6_ barrel fold are clustered with higher TM-scores (average 0.38), including CAZy defined clans GH-G, GH-L, GH-M, GH-O, GH-P, GH-Q, GH-S (total 18 families). Similarly, CAZy clans (GH-E, GH-F, GH-J) with β-propeller are also clustered with an average TM-score 0.39. The largest cluster in the heatmap contains the largest CAZy clan GH-A (28 families) and other clans (GH-D, GH-H, GH-K, GH-R, GH-T) with the (β/α)_8_ barrel fold (average TM-score 0.33).

**Figure 2:**
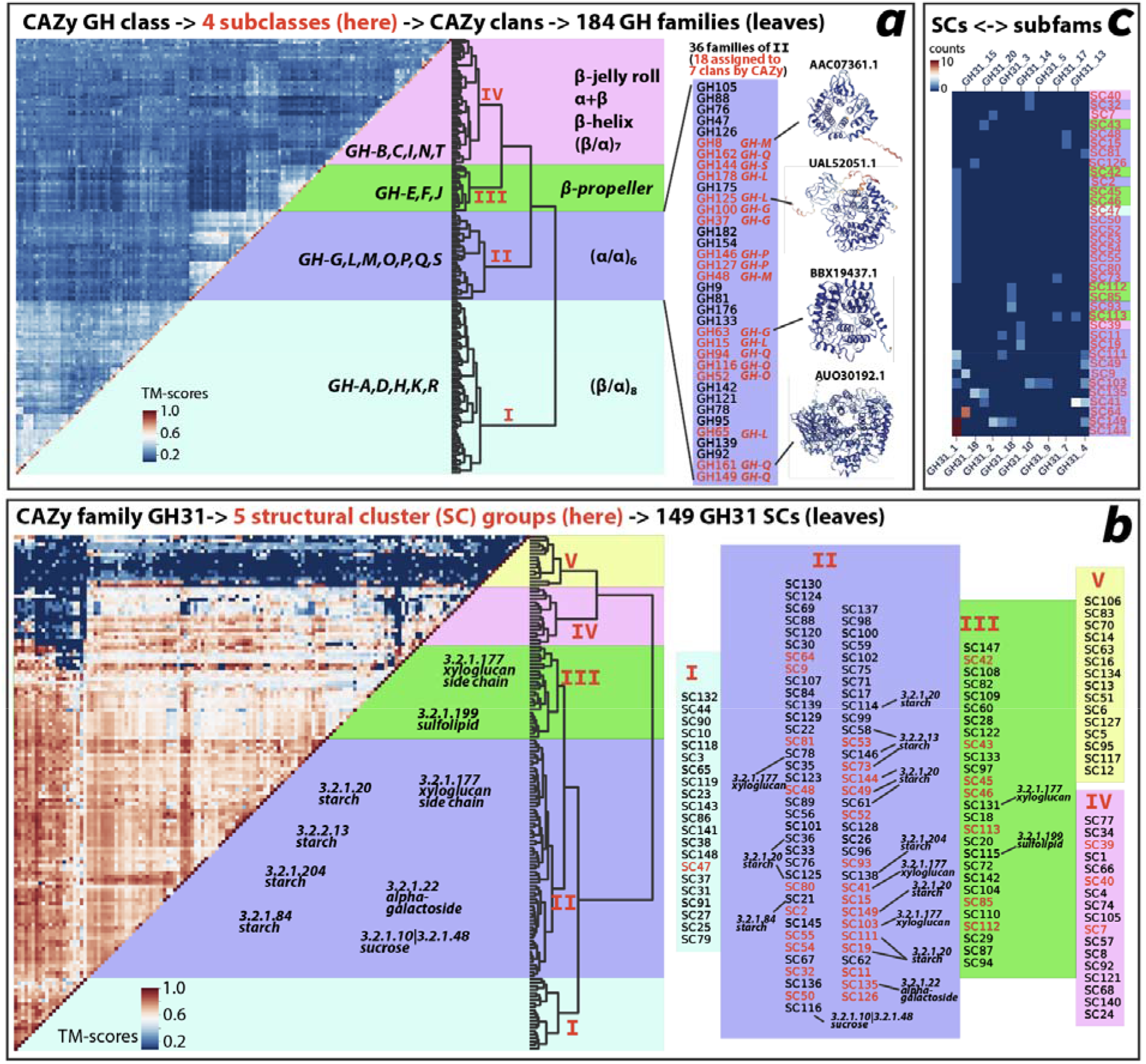
Heatmaps of structure-based hierarchical clustering of CAZymes. (**a**) Heatmap of inter-family structural comparisons of 184 families of the CAZy GH class. TM-scores (alntmscore) are calculated for each pair of GH families (see Methods). Four GH subclasses (I, II, III, IV) are defined here according to the hierarchical clustering dendrogram (leaves are 184 families). To the left of the dendrogram, 20 CAZy clans are indicated (total 76 families) in their respective subclasses. To the right of the dendrogram, structural folds of the 20 CAZy clans are indicated. Leaf names of the subclass II in (α/α)_6_ barrel fold are provided, totaling 36 families (sorted according to the dendrogram). Among the 36 families, 18 are from 7 CAZy clans (shown in red). Four example 3D structures are shown. (**b**) Heatmap of intra-family structural comparisons of 149 SCs of the CAZy GH31 family. TM-scores (alntmscore) are calculated for each pair of SCs of the GH31 family (see Methods). Five GH31 SC groups (I, II, III, IV, V) are defined here according to the hierarchical clustering dendrogram (leaves are 149 SCs). To the left of the dendrogram, EC numbers and substrates are indicated in their respective subclasses, according to contained CAZy proteins in the subclass. To the right of the dendrogram, leaf names in all the five SC groups are provided, of 149 SCs (sorted according to the dendrogram). Among the 149 SCs, 21 (19 from SC group II and 2 from SC group III) contain CAZy proteins with EC numbers and known substrates (indicated beside the SCs). Also, 36 SCs (shown in red) contain GH31 proteins that are classified by CAZy into 16 subfamilies. (**c**) Heatmap of 36 SCs mapped to 16 GH31 subfamilies according to contained CAZy ID50 protein counts. SCs are colored with different backgrounds matching those in (b).

Overall, the GH class heatmap reveals four major sub-classes (defined here as a higher level than CAZy clan, **Figure 2a**). Sub-class I contains 69 GH families with the (β/α)_8_ fold. Sub-class II contains 36 GH families with the (α/α)_6_ fold. Sub-class III contains 19 GH families with the β-propeller fold. Sub-class IV contains the rest of GH families sharing little structural similarity. All these observations also agree with our previous phylogenetic clustering of GH families using the inter-family HMM-HMM (hidden markov model) comparison method [10].

### Intra-family comparison: structural similarity guided clustering of SCs within each family

ID50 structures of each family are clustered into structural clusters (SCs) (see Methods). For example, **Figure 1b** shows that 33,423 GT2 ID50 structures are clustered into 1,317 SCs. Structures in each SCs share structural similarity and possibility also sequence similarity (but below 50% as each structure is from the nonredundant ID50 dataset, **Figure 1a**). SCs are further clustered based on pair-wise TM-scores from intra-family SC comparisons. In **Figure 2b**, we clustered 149 GH31 SCs (from 1,029 ID50 structures) into five SC groups. SC group II and III have SCs containing CAZymes with experimentally characterized ECs and substrates according to CAZy. Most of these CAZymes in SC group II degrade starch, but others degrade sucrose, xyloglucan side chain, and alpha-galactoside. CAZymes in SC group III target xyloglucan side chain and sulfolipid.

SC group is a level below family and above SC, like the CAZy defined subfamily level. We wonder how our SCs and SC groups match CAZy subfamilies. Throughout the years, CAZy has classified 35 families into subfamilies using phylogenetic classification and sequence similarity network (SSN) classification. On the CAZy website, GH31 contains 20 subfamilies (GH31_1 to GH31_20), recently created by using SSN [30]. Using the ID50 protein sequence IDs in our SCs, we found that 36 out of 149 GH31 SCs contain CAZymes from 16 CAZy GH31 subfamilies. These 36 SCs are mostly from SC groups II, III, and IV. In **Figure 2c**, we plotted the counts of CAZymes of our SCs also present in GH31 subfamilies. We observed that CAZymes from a same SC group are found in multiple GH31 subfamilies, and in most cases CAZymes from a same SC are also found in multiple GH31 subfamilies. This means that SCs or SC groups (structure-based classification) do not match CAZy subfamilies (sequence-based classification) of GH31.

### Website organization and structural similarity search web service

CAZyme structures are organized according to a seven-level hierarchical classification on our website (**Figure 3a**). From the homepage, users can access all the data via different entry options as tables or as heatmaps. Each CAZyme has its own protein page, where users can view 3D structure, sequence, functional domain, carbohydrate binding sites, and be directed to other levels (e.g., SCs, ID50 clusters, etc.) of the hierarchical classification. From the inter-family comparison pages, users can access the interactive heatmaps for all six CAZy classes, and view TM-score distribution from each family pair comparison. High quality vector images (PDF or SVG formats) can be downloaded. From the intra-family comparison pages, users can access the interactive heatmaps for 461 CAZy families. Each family page also contains a table with all SCs with EC numbers and substrates (according to contained characterized CAZy proteins).

**Figure 3:**
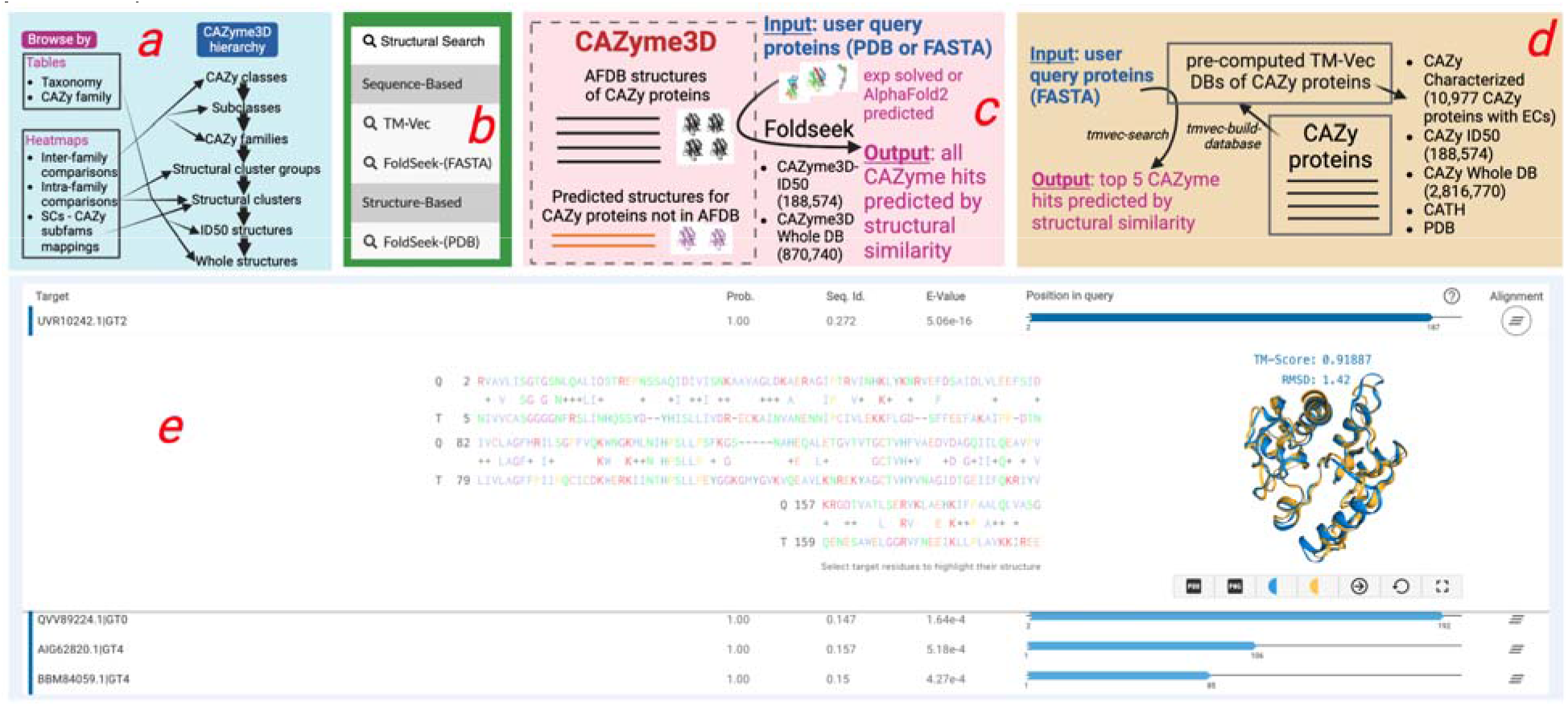
Website design of browse function and structural similarity search function. (**a**) On the right side is the hierarchical organization of data in CAZyme3D. On the left side is the list of browse options on the CAZyme3D homepage to access different data types as tables or heatmaps. (**b**) Structural similarity search drop-menu on CAZyme3D website. (**c**) Overall design of the Foldseek search page. (**d**) Overall design of the TM-Vec search page. (**e**) An example search result page (https://pro.unl.edu/CAZyme3D/results/PsIGfCfjiVsbvBsx/results.php) using a PDB protein (8FDX) as the query and CAZyme3D-ID50 database as the search DB with Foldseek structure-structure search.

Users may want to compare their own proteins against proteins in our CAZyme3D for structural similarity search. To enable this, we have constructed multiple local databases ready for TM-Vec and Foldseek searches. From the drop-down menu (**Figure 3b**) on our website, users can choose to submit FASTA sequences as query to search against the structure-aware protein embeddings of CAZymes using Foldseek (**Figure 3c**) or TM-Vec (**Figure 3d**). Users can also submit a PDB structure as query to search against all CAZyme structures in CAZyme3D using the Foldseek structure-structure search (**Figure 3c**). The results of these searches are provided as tables, where links are provided to webpages of matched CAZymes, SCs, ID50 clusters in CAZyme3D. For Foldseek structure-structure search, the HTML format output file of Foldseek (**Figure 3e**) also includes the structural alignment and the built-in visualization of superimposed structures between query and subject.

## Discussion

CAZyme3D provides data and service not existing in other databases like CAZy and dbCAN. It fills a critical gap in CAZyme research by offering 3D structure-based insights into CAZymes. Despite the presence of over 2.8 million CAZymes annotated by CAZy, only ∼10,000 experimentally solved PDB structures are available for CAZymes. CAZyme3D contains over 870,000 AlphaFold2 predicted structures for CAZymes. We did not include 3D structures for all the 2.8 million CAZymes of CAZy, as most of them failed to meet our data collection criteria (**Figure 1a**, e.g., no UniProt IDs mapped from GenBank IDs). However, all the CD-HIT representative proteins (ID50) have 3D structures included in the current release of CAZyme3D. We plan to add AlphaFold2 predicted structures for all remaining CAZy proteins in our next database release.

Using 3D structures in each CAZy class and family, we created a structure-based hierarchical classification (**Figure 3a**), including subclasses, structural cluster (SC) groups, SCs, and ID50 clusters that are newly defined here. Subclasses are used to group CAZy clans, which are only available for a minority of families in the GT and GH classes. The structural similarity-based heatmap of 184 GH families revealed four major GH subclasses (**Figure 2a**). CAZy GH clans sharing similar structural folds were accurately grouped together in the same subclasses. Heatmaps of GT, CE, AA, PL, and CBM classes can be found on their respective inter-family comparison page of the CAZyme3D website.

SC groups and SCs are defined within each CAZy family. This provides a structure-based perspective to study the intra-family CAZyme relationship, supplementing the sequence-based approaches (i.e., SSNs and phylogenies used by 35 CAZy families for subfamily classification). For example, the GH31 family contains 1,049 ID50 structures, which were first clustered into 149 SCs by Foldseek easy-cluster, and then into five major SC groups by a hierarchical clustering (**Figure 2b**). Examining ID50 CAZymes in SCs and GH31 subfamilies found that CAZymes of the same SC are present in multiple CAZy subfamilies, and *vice versa*. The intra-family comparison heatmaps are available for all CAZy families on our website.

## Code and data availability

All data of CAZyme3D can be downloaded https://pro.unl.edu/CAZyme3D/. CAZyme3D will be updated yearly to include AlphaFold2 predicted 3D structures of new CAZymes.

## CRediT authorship contribution statement

**N.R. Siva Shanmugam**: Writing – review & editing, Writing – original draft, Visualization, Database construction, Methodology, Formal analysis, Data curation, Conceptualization.

**Yanbin Yin:** Writing – review & editing, Supervision, Conceptualization.

## Declaration of competing interest

The authors declare no competing interests.

## Acknowledgments

This work was partially completed utilizing the Holland Computing Center of the University of Nebraska-Lincoln.

## Funding

This work was supported by the U.S. National Institutes of Health (NIH) awards [R01GM140370] and [R21AI171952], U.S. Department of Agriculture (USDA) award [58-8042-9-089]. Funding for open access charge: NIH award [R01GM140370]. This work was also partially completed utilizing the Holland Computing Center of the University of Nebraska, which receives support from the Nebraska Research Initiative.

## Notes

https://pro.unl.edu/CAZyme3D/

### Competing Interest Statement

The authors have declared no competing interest.

